# Fundamentals on the Kinetic and Thermodynamic Analysis of Oligonucleotide DNA Hybridization by Surface Plasmon Resonance: A Guide for HIF1α Antisense Design

**DOI:** 10.64898/2026.08.10.743984

**Authors:** Stephen Cornwell, Frank Podlaski, Kenny Wong, Brian McKittrick, Jae-Hun Kim, William T. Windsor

**Author notes:** These authors contributed equally to this work and are to be considered primary authors (Stephen Cornwell, Frank Podlaski). Research & Development, Vaccines, Pfizer Inc., 401 N. Middletown Road, Pearl River, NY 10965. Drew University, 36 Madison Ave, Madison, NJ 0794O. JK Pharm Consulting LLC, 242 Denman Rd, Cranford, NJ 07016. <u>E-mail Addresses:</u> Stephen Cornwell, Frank Podlaski, Kenny Wong, Brian McKittrick, Jae-Hun Kim. Corresponding Author: William T. Windsor, Department of Chemistry and Chemical Biology, Stevens Institute of Technology, 1 Castle Point Terrace, Hoboken, NJ, 07030, USA.

## Abstract

Antisense oligonucleotides (ASO) are nucleotide polymers that hybridize to sense strands and have been successful in treating a variety of diseases. A wide range of strategies have been investigated to optimize and develop ASO for clinical studies. A key objective for this study was to provide an overview of the range of detailed data that can be obtained and provide an updated method review on how to design surface plasmon resonance (SPR) kinetic experiments for DNA oligonucleotide hybridization studies that can also be applied to other ASO including peptide nucleic acids (PNA). We describe many lessons learned from published literature and provide a state-of-the-art strategy and methods for generating not only kinetic but also thermodynamic characterizations of oligonucleotide hybridization. In this study we have performed an SPR kinetic and thermodynamic analysis for the hybridization of HIF1α antisense DNA strands to its immobilized Intron2-Exon3 splice site sense DNA strand to provide insight, in general, on the optimal length and insight into optimal design of DNA ASOs. We provide a process on how to design experiments to: 1.) obtain oligonucleotide-length dependent kinetics, 2.) analyze reactions to obtain association and dissociation rate kinetics (k_a_, k_d_), assess if hybridization follows a 2-state model and to obtain kinetic dissociation constants (K_d_), 3.) perform temperature-dependent hybridization kinetics to obtain thermodynamic values (ΔH°, ΔS° and ΔG°) that can give insight into the molecular interactions driving hybridization, 4.) compare experimental thermodynamic values to values derived from nearest-neighbor prediction models to identify atypical reactions and importantly 5.) enable calculations to predict oligomer hybridization affinity at the physiological 37 °C temperature to asses if the design of the oligomer will have the required cellular activity for a therapeutic effect. The strategy and results presented throughout the paper are compared to previous SPR reports and suggestions made to optimize kinetic studies.

## Introduction

Antisense oligonucleotides (ASO) are nucleotide polymers that bind complementary sequences of RNA or DNA oligonucleotides. ASO have been successful in treating a variety of diseases including muscular, neurological, immunology, vision and metabolic. ^1,2^ The mechanism of action for these disease modifiers includes the stimulation of RNA degradation by RNAse H1, modification of pre-mRNA splicing to mature mRNA and immune stimulation by recruiting proteins to regulator sites^2,3^. The successful clinical application of the ASO strategy has been mediated by the design of novel modification to the oligonucleotides to provide enhanced metabolic stability, increased affinity to targeted nucleotide sequences and cell permeability^4,5^. One of the first methods to prevent hydrolysis of these typically short oligonucleotides (10-20 base pairs) is to replace the phosphate backbone with a peptide nucleic acid (PNA) backbone which utilizes a modified N-(2-aminoethyl)glycine peptide backbone and attached nucleotide bases without the sugar component^4,6–8^. While the PNA design retains the ability to have excellent nucleotide hybridization properties and is resistant to hydrolysis it has poor cell permeability and requires high concentrations in both cell-based and *in vivo* models to observe effects on target levels and function. To improve cell-permeability, several strategies have been employed including formulating/encapsulating the ASO with phospholipids as well as introducing additional chemical modifications such as adding positively charged groups like lysine amino acids, to the 5’ (N-terminus)/3’ (C-terminus) ends of the synthetic oligonucleotide^4,5,9,10^. Another strategy to improve cell-permeability is to introduce positive charges directly on the ASO nucleotide bases^11^. In a previous publication we described how to perform SPR-based hybridization kinetics and thermodynamics of a single-strand PNA oligonucleotides containing a cationic/amino-alkyl modified bases to DNA oligonucleotides^12^.

Selection of optimal synthetic single strand ASO oligonucleotides requires studying many different molecules in *in vitro* assays that measure affinity, stability, cellular response, in addition to *in vivo* models. Cell-based assays and *in vivo* models vary greatly depending on the intended target and desired disease outcome. However, the analytical methods for characterizing the thermodynamic, kinetic, and thermal stability of oligonucleotide hybridization are performed by only a few different types of methods. Over the course of the last 40 years one of the primary methods for assessing the affinity of ASO hybridization with target DNA or RNA single strands is to measure the thermal stability of ASO/oligonucleotide double strands. Such studies can provide a thermodynamic characterization of traditional DNA, RNA or oligo modified ASO double strands providing estimates of enthalpy (ΔH°), entropy (ΔS°) and free energy (ΔG°) of hybridization^13^. Experimentally derived thermodynamic parameters have led to the development of predictive models such as the nearest-neighbor model from which the thermodynamic parameters can be calculated solely on a double strand nucleotide sequence which informs on the type of Watson and Crick base-pair hydrogen bonds and stacking energetics between two bases^14–19^. Kinetic characterization of hybridization rates is also of value to provide insight into the mechanism of association which, for short oligonucleotide strands, can identify unexpected non-two state binding issues for association potentially due to steric hindrances such as unexpected hairpins^20–23^. In addition, kinetic studies measuring the hybridization rates of association and dissociation to provide binding constants versus temperature can provide similar thermodynamic parameters as the nearest-neighbor method using the van’t Hoff Plot.

For the last ~ 25 years a common method for kinetic analysis of hybridization rates has been to use some derivation of immobilizing a target DNA sequence and measuring a signal upon binding the complementary single strand oligonucleotide. The techniques and signals most often used include time-dependent fluorescence enhancement or quenching and surface plasmon resonance (SPR) with and without fluorescence^24–29^. In this current report we have used SPR to measure the kinetics of short 10 to 13-mer single strand DNA (ssDNA) oligonucleotides to a conserved biotin-immobilized complementary 17-mer ssDNA. Throughout the report we describe methods that should be applied when performing SPR ASO hybridization studies and compare these methods to studies that were not optimized in earlier reports. The origin of the 17-mer target is from the HIF1α Intron2-Exon3 “3’-splice site”^30^. HIF1α is a target for inhibition due to its role in promoting cancer growth by tumors that are in hypoxic environments^31,32^. Our ssDNA ASOs were designed to bind across the splice site similar to a previously described series of PNA ASOs that target the same splice site and for which we have demonstrated robust inhibition of HIF1α mRNA and protein expression and inhibition of cancer cell growth^33^. Our current experiments were designed to study the fundamental molecular kinetic and thermodynamics that promotes hybridization which might provide insight into the design of PNA oligonucleotides at the splice site. From length-dependent and temperature-dependent kinetics of DNA-DNA hybridization it is apparent that dimerization for the short ASO strands is consistent with a two-state model indicating the selected sequences do not form undesirable hairpins, or secondary structures that would hinder binding and lead to non-2-state binding ^35^. In addition, the biomolecular association rate appears to be mostly independent of nucleotide length for 10 to 13mers while the dissociation rate decreases greatly per added base-pair. Thus, the affinity is driven nearly exclusively by the dissociation rate, likely due to the additional energetics stemming from sequential additions of Watson and Crick base-pair hydrogen bonds and base-base stacking energetics. From temperature-dependent kinetic studies, thermodynamic analysis provided ΔH°, and ΔS° to which we were able to compare to predicted values using nearest-neighbor models that accounts for specific NaCl concentrations^17^. The excellent comparison of kinetic-derived thermodynamic parameters to the predicted nearest-neighbor models indicates the modified methods we used in the SPR studies provide a reliable and alternative method for generating important thermodynamic parameters for short oligonucleotide hybridization studies.

## Material and Methods

### Oligonucleotides

Sense and antisense single strand DNA oligonucleotides were synthesized by Integrated DNA Technologies (Coralville, IA). Immobilized sense strands were labeled with biotin at the 5’ end. All oligonucleotides were HPLC purified, confirmed by mass spectroscopy and suspended in deionized water to make a working 500 µM stock.

### Surface plasmon resonance kinetics and analysis

All SPR experiments were carried out using a Sartorius Pioneer FE (Sartorius BioAnalytical Instruments, Inc, Fremont, CA) and streptavidin (SADH) biosensors from FortéBio (Part #19-0130). All experiments, unless otherwise specified, were carried out at 20 °C. Experiments utilized a consistent running buffer composed of HBS-T (HEPES-buffered saline, 10 mM HEPES, 150 mM NaCl, pH 7.4 with 0.005% Tween-20). To prevent buffer mismatch response upon sample injection, all samples were diluted from a concentrated deionized water stock into this running buffer such that the difference in buffer composition was negligible. All buffers were filtered using 0.45μm pore filters prior to use.

All experiments began with priming the system with running buffer three times, followed by installation of the streptavidin (SADH) sensor chip. All three flow cells were checked for alignment of SPR angle. The sensor was conditioned using 50 mM NaOH in deionized water over all flow cells at 50 µL/min for 1 minute. The biotinylated 17-mer sense strand was immobilized to the surface of flow cell 1 (FC1) by manual injection of 100 nM solution at 10 µL/min until a capture response of approximately 100-150 RU was seen. 100 nM of a mutated 17-mer sense ssDNA (scrambled ASO sequence) was immobilized on flow cell 3 (FC3) at the same response level (100-150 RU) to ensure specific binding was observed on FC1. FC2 was used as a reference cell with no target immobilized. To block any potential unreacted streptavidin sites, 20 µM biotin (Aldrich, CAS #58-85-5) in running buffer was flowed over all cells (FC1, FC2, and FC3). Buffer solution was injected over all flow cells at 50 µL/min for 30 seconds twice prior to running any assay to ensure that responses were consistent, and that signal returned to baseline following injections.

Dilution series of all analytes to be tested (AS-10 through AS-13) were made such that two-fold, five-point concentration ranges were made in running buffer unless otherwise specified. The appropriate concentration range for each analyte was determined via initial manual injection experiments (data not shown). Experiments were run in succession on the same flow cell at 50 µL/min, with 180 seconds for analyte injection and approximately 300 seconds for dissociation (unless otherwise noted). Each injection cycle was followed by injection of 10 mM HCl for 30 or 60 seconds to regenerate the surface as was determined from trials of manual injections to re-establish the original baseline. Each experiment conducted was performed at least in duplicate, and reported analytical data represents an average of these experimental results. Useful experimental details for performing PNA SPR kinetics can be found in our previous publications^12^.

All data was obtained using Pioneer Instrument Software version 4.3.1, build 32. Analysis of sensograms to obtain the RU binding amplitude, on rates, off rates, and affinity measurements was performed using Qdat^®^ Data Analysis Tool, version 4.3.1 build 2. All curves were fit to a bimolecular reaction using the global non-linear least squares regression Analysis Tool, first for only the off rate, then the off rate was fixed once to help solve for the on rate before finally fitting both the on and off rates and RUmax.

## Results and Discussion

### 1. Kinetic hybridization of short HIF1α antisense ssDNA

ASO hybridization studies were performed using oligonucleotide sequences from the pre-mRNA HIF1α Intron2-Exon3 “3’ splice site” as described in the PNA study by Chung et. al., (2018) ^33^. The binding kinetics of four complementary antisense oligonucleotides (ASO) to the sense strand were measured by surface plasmon resonance using a streptavidin SADH sensor chip to which the corresponding 5’-modified biotin-17mer HIF1α DNA sense strand (b-17mer-Sens) was immobilized (Table 1). The four ASO were designed to span across the 3’ splice site, but each ASO increased by 1 nucleotide by alternating the addition from the 3’ to the 5’ site yielding strands AS-10 (10mer), AS-11 (11mer), AS-12 (12mer) and AS-13 (13mer). At the beginning of this study, we had synthesized each of the 10mer to 17mer antisense strands that would complement the biotin-17mer sense strand, but it became apparent that the dissociation rate for oligomers longer than 13mer at 20 °C had a dissociation rate that was too slow (k_d_ < 4 ×10^−4^ s^−1^, t_1/2_ ~ 2 hrs, data not shown) for accurate kinetic analysis. The very slow dissociation rate also prevented accurate measurements of the association rate (k_a_) and affinity (K_d_), for the 14 to 17mers, therefore, these studies only used the 10 – 13mer sequences.

**Table 1.**
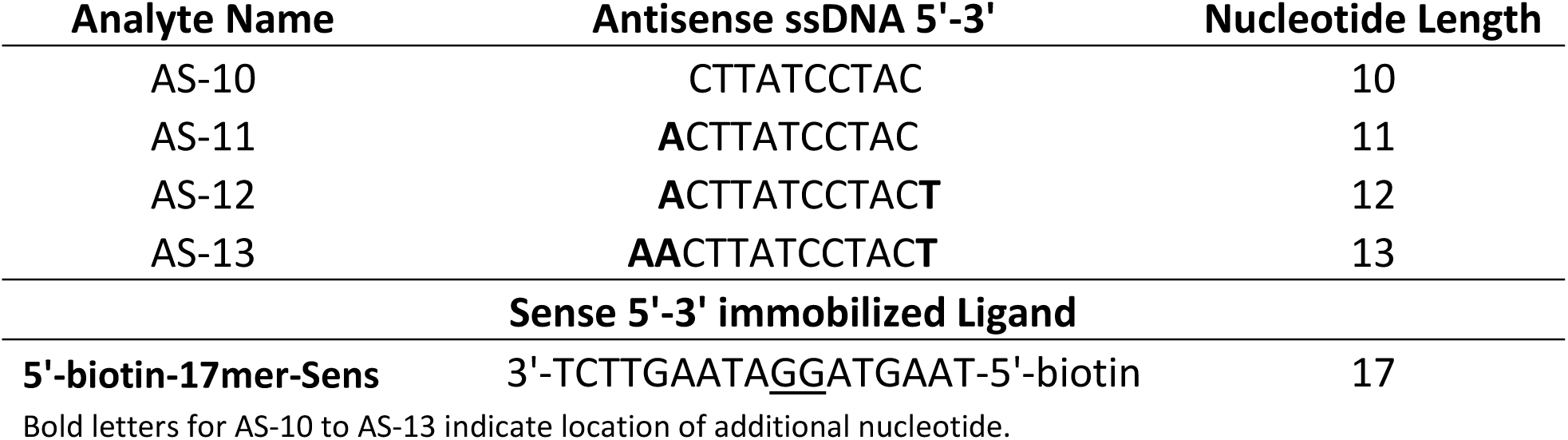
HIF-1 Antisense Sequences for Binding Studies.

Kinetic studies were performed for each ASO at 20 °C, 25 °C and 30 °C and Figure 1 shows the typical binding responses observed for AS-10, AS-11, AS-12 and AS-13 at 25 °C. These data show that there is a well-defined concentration dependence of binding for each ASO leading to a maximum response consistent with near saturation of the immobilized b-17mer-Sens target. One of the key experimental designs for these SPR studies was to minimize the loading of the b-17mer-Sens strand to prevent a commonly observed mass transport diffusion artifact that inhibits an accurate analysis of rate constants^36,37^. In these studies, the total load of b-17mer-Sens onto the sensor chip was kept at a minimum where the mass loaded to the sensor provided only ~100-120 RU. This low load would then minimize the mass transport issue and would produce a maximum signal upon full binding of the ASO strands to the b-17mer-Sens of only ~70-80 RU which is exactly what is observed in Figure 1. Literature search of previous SPR studies find that many of the early studies did not use low immobilized target densities (eg., RU Loads ~ 3,000 and RUmax ~ 2,000) which may have affected the final kinetic analysis and lead to underestimating values^38^. Confirmation that the biomolecular binding is a two-state event and not influenced by mass transport diffusion is supported by: 1.) the very good bimolecular non-linear least squared fit (dashed line in Figure 1) and 2.) no change in shape of the binding curve at different flow rates (not shown). Comparison of the 10mer AS-10 curves to the longer 13mer AS-13 curves shows a clear decrease in dissociation rate as would be expected due to the stabilization energetics through the addition of 3 more base pairs. Increasing the temperature to 30 °C increased the dissociation rate for both ASO strands which is consistent with the elevated thermal energy weakening both hydrogen bonds and base stacking energetics (data not shown).

**Figure 1.**
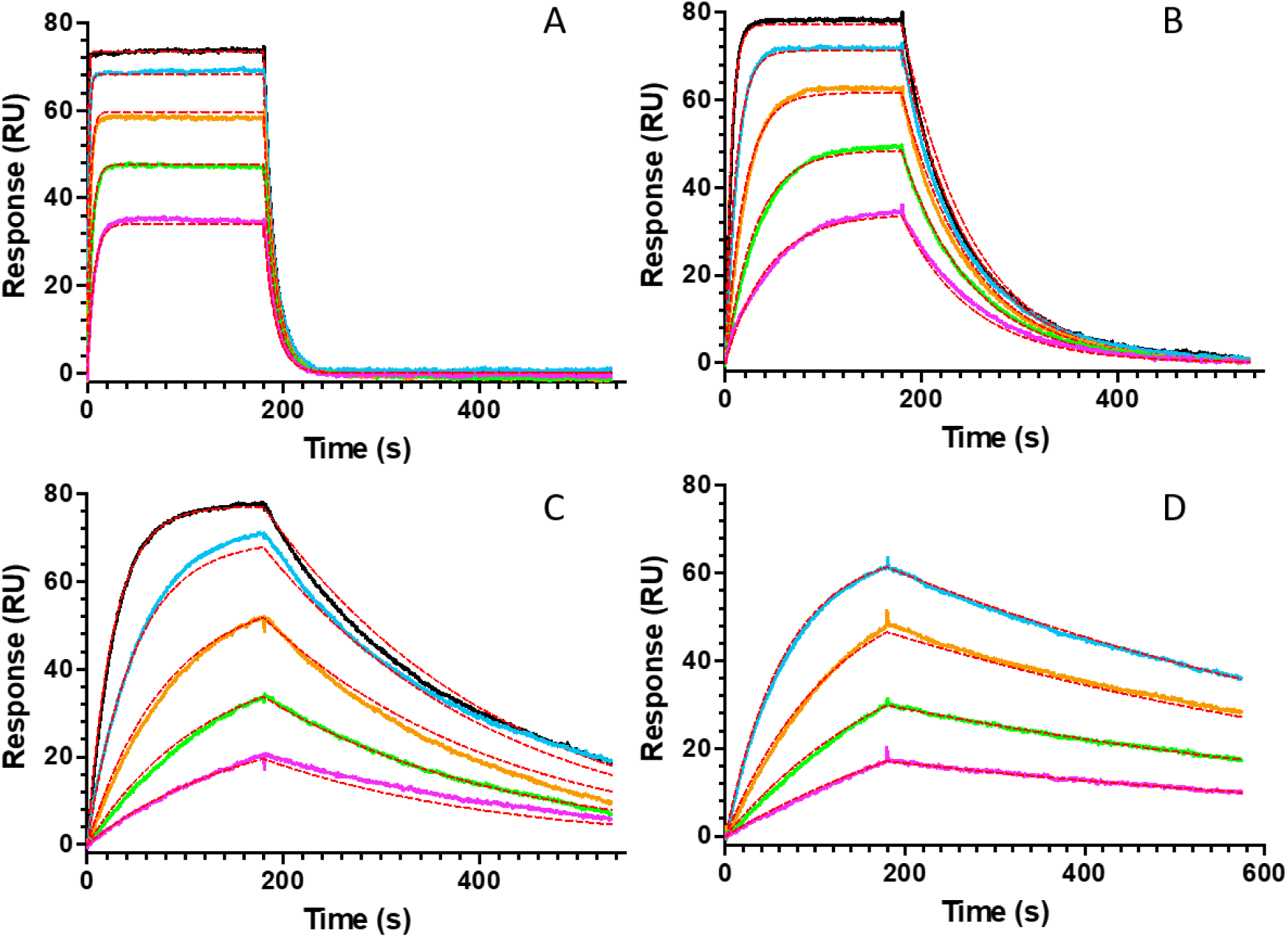
SPR kinetic profiles for antisense hybridization at 25 °C. (A), (B), (C) and (D)are AS-10, AS-11, AS-12 and AS_13 kinetic curves at 25 °C, respectively. Concentrations of antisense strand are 2-fold dilutions from 6.25 nM to 100 nM except for AS-13 (6.25 nM to 50 nM). The association reactions start from time zero to 190 sec followed by buffer wash and ASO dissociation. Red dashed lines are NLLSQ fit to the global fit.

The full kinetic analysis for AS-10, AS-11, AS-12 and AS-13 at 20 °C are presented in Table 2. The k_a_ value for AS-10 is 2.8 x 10^5^ M^−1^s^−1^ and increases only ~ 2-fold for AS-13 despite the addition of 3 more nucleotides (Figure 2A). The relative insensitivity of association rates is consistent with intrinsic diffusion rates of association that are not affected by small changes in molecular weight or the effect of increasing length on polymer geometrical shape or radius of gyration^39^. The value of the observed k_a_ rates is consistent with that observed by a wide range of kinetic studies performed with short nucleotides (9-20 nt)^24,40–42^.

**Figure 2A.**
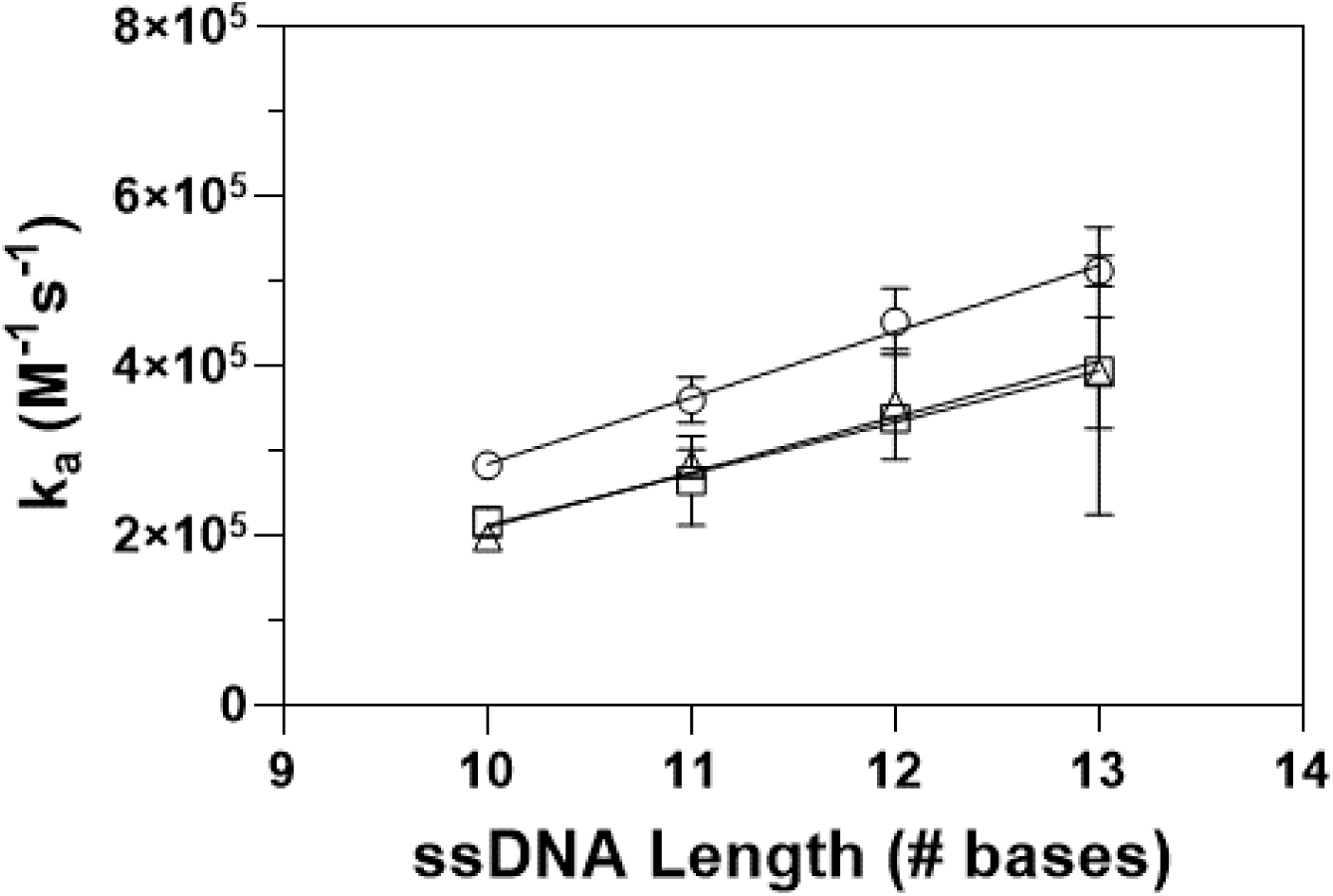
Length and Temperature-dependence for the association rate constant of the AS-10, AS-11, AS-12, AS-13 oligonucleotide hybridization. Circle: 20 °C, Square: 25 °C, Triangle: 30 °C.

**Table 2.** Kinetic Analysis for Hybridization.

| Temperature | Analyte | $k_a$<br>($10^5 \text{ M}^{-1}\text{s}^{-1}$) | $k_d$<br>($10^{-4} \text{ s}^{-1}$) | $K_d$ (nM) | $t_{1/2}$ (m) |
| --- | --- | --- | --- | --- | --- |
| 20 °C | OP3-10 | $2.83 \pm 0.13$ | $151 \pm 29.0$ | $53.7 \pm 12.7$ | 0.78 |
| 20 °C | OP3-11 | $3.60 \pm 0.27$ | $20.7 \pm 0.74$ | $5.75 \pm 0.22$ | 5.59 |
| 20 °C | OP3-12 | $4.52 \pm 0.39$ | $7.04 \pm 0.42$ | $1.57 \pm 0.23$ | 16.43 |
| 20 °C | OP3-13 | $5.12 \pm 0.18$ | $1.01 \pm 0.03$ | $0.20 \pm 0.01$ | 114.48 |
| 25 °C | OP3-10 | $2.17 \pm 0.18$ | $816 \pm 54$ | $380 \pm 57$ | 0.14 |
| 25 °C | OP3-11 | $2.65 \pm 0.53$ | $134 \pm 9.0$ | $51 \pm 7$ | 0.87 |
| 25 °C | OP3-12 | $3.38 \pm 0.08$ | $42.1 \pm 1.6$ | $12 \pm 0.2$ | 2.75 |
| 25 °C | OP3-13 | $3.94 \pm 1.70$ | $13.1 \pm 0.08$ | $3.7 \pm 1.8$ | 8.87 |
| 30 °C | OP3-10 | $1.98 \pm 0.07$ | $3,970 \pm 83$ | $2,000 \pm 29$ | 0.03 |
| 30 °C | OP3-11 | $2.85 \pm 0.17$ | $858 \pm 74$ | $302 \pm 43$ | 0.14 |
| 30 °C | OP3-12 | $3.55 \pm 0.65$ | $256 \pm 22$ | $73 \pm 7.0$ | 0.45 |
| 30 °C | OP3-13 | $3.92 \pm 0.66$ | $51.0 \pm 3.7$ | $13 \pm 3.2$ | 2.27 |

The k_d_ rates, however, decrease greatly as each additional nucleotide is added. AS-10 has a relatively fast off rate of 150 x 10^−4^ s^−1^ (0.015 s^−1^, or half-life: t_1/2_ = 0.78 min) while AS-13 has an off rate of 1.0 x 10^−4^ s^−1^ (t_1/2_ = 1.9 hrs) at 20 °C (Table 2). The addition of 3 nucleotide base pairs decreases the k_d_ rate exponentially by 150-fold or approximately 5.3-fold per base pair (Figure 2B). The hybridization equilibrium dissociation constant (K_d_) can be calculated from the ratio of k_d_/k_a_ and is presented in Table 2 for 20 °C. AS-10 10mer has a K_d_ of 54 nM and the K_d_ decreases to 0.20 nM for the 13mer, AS-13. This approximately 270-fold enhancement in affinity for AS-13 comes primarily from the 150-fold decrease in k_d_ rate and appears to be an exponential function of the sequence length (Table 2, Figure 2B). Similar observation in the k_a_, k_d_ and K_d_ values by length have been reported previously.^42–44^ Thus, the affinity of short oligomers is primarily a function of the number of nucleotides in the sequence, the number of base pair hydrogen bonds and the number, as well as type of stacking base pair interactions which stabilize the duplex decreasing the rate of dissociation.

**Figure 2B.**
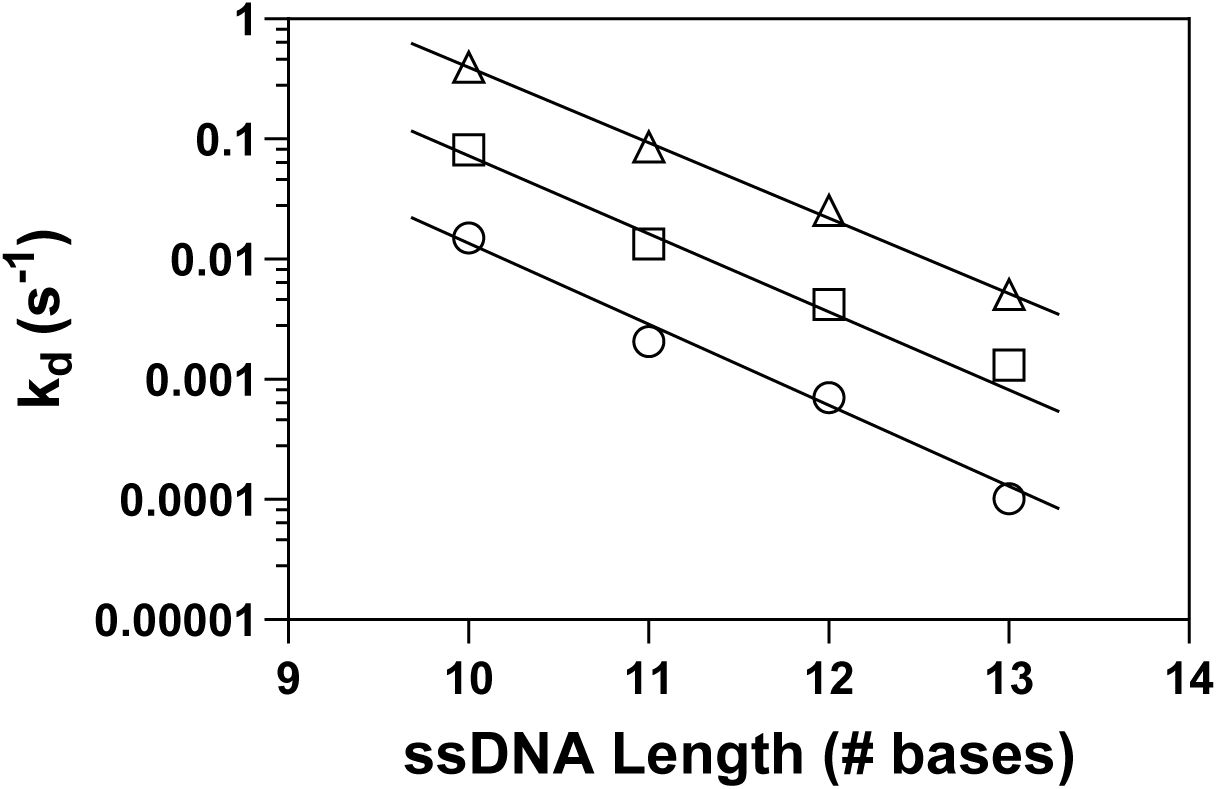
Length-dependence for the dissociation rate constant of the AS-10, AS-11, AS-12, AS-13 oligonucleotide hybridization. Circle: 20 °C, Square: 25 °C, Triangle: 30 °C.

The nearly linear k_d_ dependence on sequence length seen in Figure 2B may allow predictions of dissociation rates for other short lengths not studied, assuming a similar association rate for each. Some predicted k_d_ values (based on Fig 2B) are: 5mer, 43 s^−1^ (t_1/2_ 1.6 x 10^−2^ s); 15mer, 4.3 x 10^−6^ s^−1^ (t_1/2_ 44 hr); 17mer 1.7 x 10^−7^ s^−1^ (t_1/2_ 1,100 hr) and 20mer, 1.4 x 10^−9^ s^−1^ (t_1/2_ 140,000 hr) at 20 °C. Assuming an invariant value of k_a_ of 2 x 10^5^ M^−1^s^−1^ from our study the corresponding K_d_ for a 17mer ASO would be ~ 0.01 nM. Importantly, these k_d_ dissociation rate results suggests that many of the previously published DNA and PNA kinetic studies performed studying hybridization kinetics for oligomers > 14mer would have to design the studies to measure off rates on the order of days. These very slow dissociation rates, however, in general were not observed or reported and it is not known why many other studies had faster dissociation rates. Indeed, our initial hybridization studies planned to measure up to 17mers based on published literature, however, it became apparent that only dissociation rates for oligomers less than 14mer could be recorded accurately by SPR at 20 °C.

Performing temperature-dependent kinetic binding studies provides insight into the thermal stability of hybridization and identifies effects on the association and dissociation rate constants which can provide information on molecular interactions that drive the dimerization event. Inspection of the k_a_ values (Table 2, Figure 2A) for each of the ASO with respect to base length and temperature shows that the k_a_ rate for each ASO is marginally sensitive to an increase in 10 °C. This relative insensitivity for the on rate is a common observation reflecting that elevated solvent energetics is not strongly dependent on diffusion and bimolecular recognition/initiation at these moderate temperatures^34,44^. However, the dissociation rate k_d_ for each ASO increases exponentially with temperature and is ~25-50-fold faster for each ASO at 30 °C versus 20 °C (Figure 2C). The thermally sensitive molecular interactions that lead to increased dissociation rates are due primarily to the weakening of base pair hydrogen bonds, dipole stacking interactions and increased fraying of the 5’ and 3’ ends. The thermal effects on k_a_ and k_d_ are consistent with other DNA-DNA and PNA-DNA hybridization systems^44–46^.

**Figure 2C.**
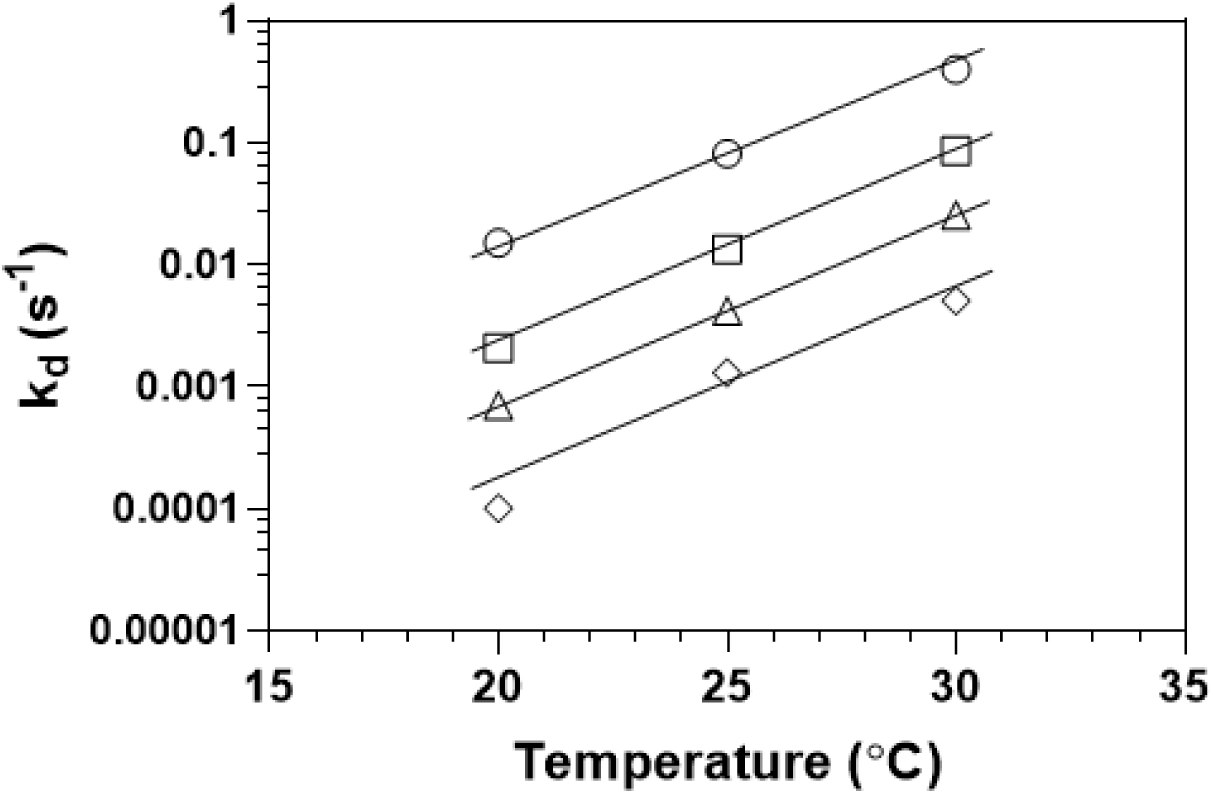
Temperature-dependence for the dissociation rate constant of the AS-10, AS-11, AS-12, AS-13 oligonucleotide hybridization. Circle: AS-10, Square AS-11, Triangle: AS-12, Diamond: AS-13.

### 2. Hybridization Thermodynamic Analysis

The equilibrium dissociation constants (K_d_) calculated from the SPR hybridization rate constants for experiments performed at 20 °C, 25 °C and 30 °C are presented in Table 2 for each of the ASO. The AS-10 oligomer K_d_ increases from 53 nM to 2,000 nM from 20 °C to 30 °C. The affinity decrease (K_d_ increase) with increasing temperature is consistent with the fold increase in the dissociation rate k_d_. The average fold change in K_d_ for all 4 ASO from 20 °C to 30 °C is ~ 45-fold (AS-10 to AS-13: 37, 52, 46, 65-fold, respectively). While these temperature-dependent changes in kinetic rates and affinity can provide insight into the molecular interactions mediating the hybridization (e.g., 3-D diffusion, base pair formation/initiation and hybridization zippering and confirmation of 2-state bimolecular binding)^34^ information regarding the energetic contributions toward hybridization can be provided by calculating the free energy of binding, ΔG°, as well as the enthalpy, ΔH° and entropy of binding ΔS°, with the relationship ΔG° = ΔH° - TΔS° describing the full thermodynamic relationship. The free energy is also a function of the equilibrium association binding constant (K_a_), ΔG° = - RTln(K_a_), where the association binding constant K_a_ can be calculated from the kinetic derived affinity 1/K_d_, R is the universal gas constant and T is the temperature in Kelvin. Table 3 shows the kinetic-derived ΔG° for each of the ASO at the three temperatures.

**Table 3:** Kinetic and Nearest Neighbor Thermodynamic Analysis for Hybridization.

| Kinetic Thermodynamic Analysis |  |  | 20° C |  | 25° C |  | 30° C |  |
| --- | --- | --- | --- | --- | --- | --- | --- | --- |
| Analyte | $\Delta H$ (kJ/mol) | $\Delta S$ (J/mol K) | $-T\Delta S$ (kJ/mol) | $\Delta G$ (kJ/mol) | $-T\Delta S$ (kJ/mol) | $\Delta G$ (kJ/mol) | $-T\Delta S$ (kJ/mol) | $\Delta G$ (kJ/mol) |
| AS-10 | $-267 \pm 11$ | $-777 \pm 39$ | 227 | $-41 \pm 0.6$ | 231 | $-37 \pm 0.4$ | 235 | $-33 \pm 0.1$ |
| AS-11 | $-292 \pm 10$ | $-840 \pm 33$ | 246 | $-46 \pm 0.1$ | 251 | $-42 \pm 0.8$ | 255 | $-38 \pm 0.4$ |
| AS-12 | $-283 \pm 8$ | $-800 \pm 28$ | 235 | $-49 \pm 0.4$ | 239 | $-45 \pm 0.6$ | 243 | $-41 \pm 0.2$ |
| AS-13 | $-310 \pm 37$ | $-874 \pm 125$ | 256 | $-54 \pm 0.2$ | 262 | $-48 \pm 1.8$ | 265 | $-46 \pm 0.6$ |
| **Nearest Neighbor Thermodynamic Analysis** |  |  |  |  |  |  |  |  |
| AS-10 | -289 | -846 | 248 | -41 | 252 | -37 | 256 | -33 |
| AS-11 | -315 | -913 | 268 | -47 | 272 | -43 | 277 | -38 |
| AS-12 | -338 | -974 | 286 | -52 | 290 | -47 | 295 | -42 |
| AS-13 | -370 | -1066 | 312 | -57 | 318 | -52 | 323 | -46 |

A full thermodynamic characterization of the hybridization reactions can be determined using the van’t Hoff Plot that linearizes the relationship of K_a_ to inverse temperature giving ln(K_a_) = - ΔH°/RT + ΔS°/R which enables the determination of both the enthalpy and entropy of binding from the temperature-dependent SPR kinetic experiments. Figure 3 is the van’t Hoff plot and graphically indicates a linear relationship between ln(K_a_) and the inverse temperature which because the response is linear permits calculations of the enthalpy, ΔH°, and entropy, ΔS° of hybridization (assuming the heat capacity, ΔC_p_°, of binding is small over the temperature range). Table 3 summarizes the results for each of the ASO. The negative total ΔH° values for AS-10, AS-11, AS-12, and AS-13 are −267 kJ/mol, −293 kJ/mol, −284 kJ/mol and −310 kJ/mol respectively and when normalized by number of base pairs per heterodimer gives −26.7 kJ/mol, −26.6 kJ/mol, −23.7 kJ/mol and −23.8 kJ/mol per base pair, respectively. These favorable enthalpies per base pair values represent the stabilizing energetics that originate from the 2 or 3 Watson and Crick Hydrogen bonds for A:T and G:C pairs, respectively, as well as the base pair / base pair dipole stacking energetics and are similar to values reported by other groups^14,15,19^.

**Figure 3.**
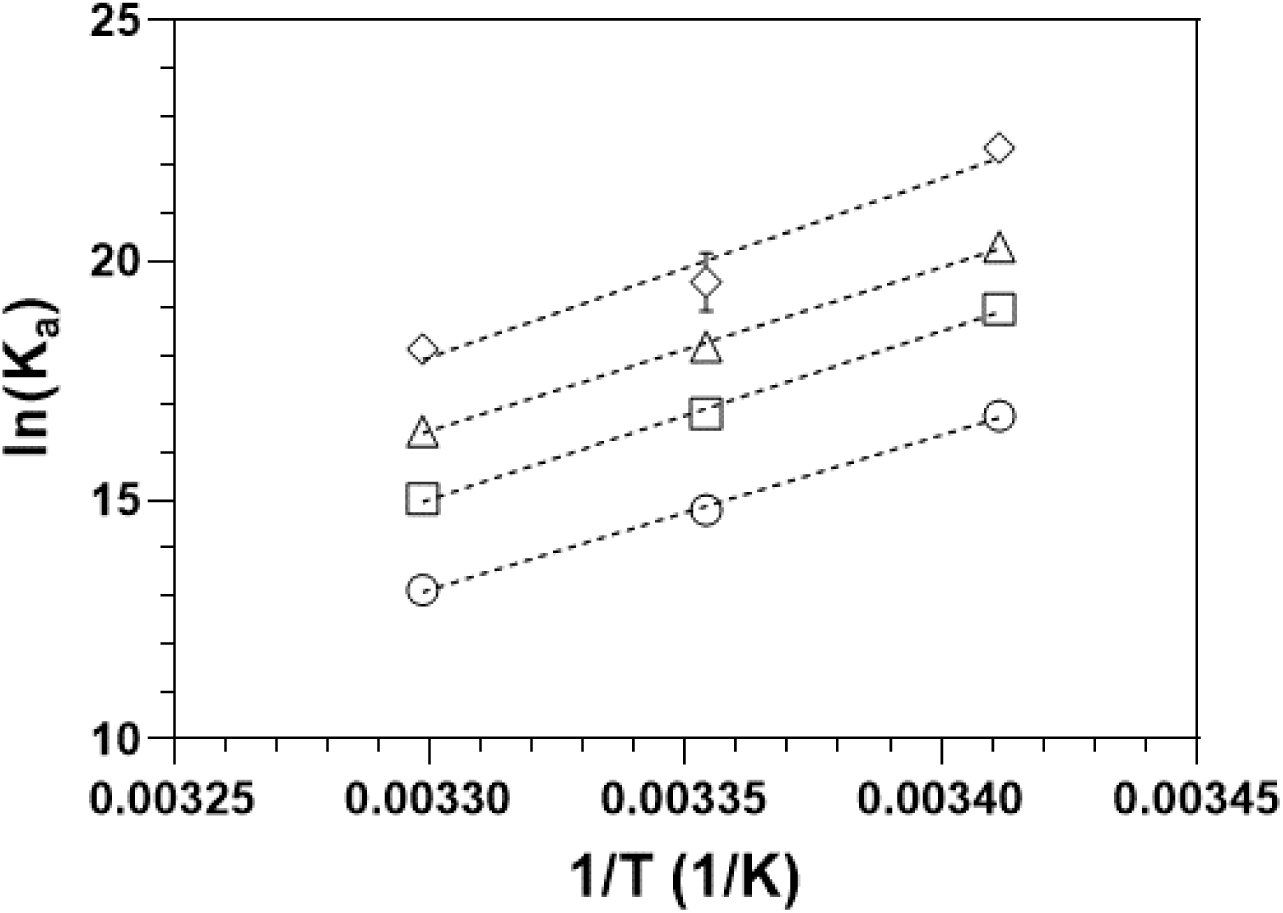
Van’t Hoff plot to calculate enthalpy and entropy values from the ASO hybridization kinetic studies. Circle: AS-10, Square: AS-11, Triangle: AS-12, Diamond: AS-13. Equilibrium association constant (K_a_) derived from the K_d_ value in Table 2. Enthalpy values for each ASO derived from the linear plots are reported in Table 3.

The entropy contribution to the free energy, −TΔS°, for AS-10 at 20 °C, 25 °C and 30 °C shown in Table 3 are +227 kJ/mol, +231 kJ/mol and +235 kJ/mol. These entropy terms are unfavorable energetically and likely due primarily to the induced conformational constraint of the unstructured flexible single strand upon formation of the constrained double strand helical structure as-well-as solvent effects. Similar unfavorable entropy values are seen for the other ASO going from 20 °C to 30 °C^14,15,19^. In addition, it is clear the magnitude of the −TΔS° term does not increase greatly going from AS-10 to AS-13 suggesting that 3 additional nucleotides do not introduce a large entropic penalty. These temperature studies demonstrate that the overall free energy of binding is driven by the favorable enthalpy energetics upon hybridization. Figure 4 shows that the ΔG° of binding is linear with increasing base pair length and indicates each additional base pair from 10 to 13 promotes on average −3.8 kJ/mol per base pair stabilization at 20 °C, 25 °C and 30 °C. The average change in ΔG° per degree Celsius for each ASO is +0.82 kJ/mol/°C.

**Figure 4.**
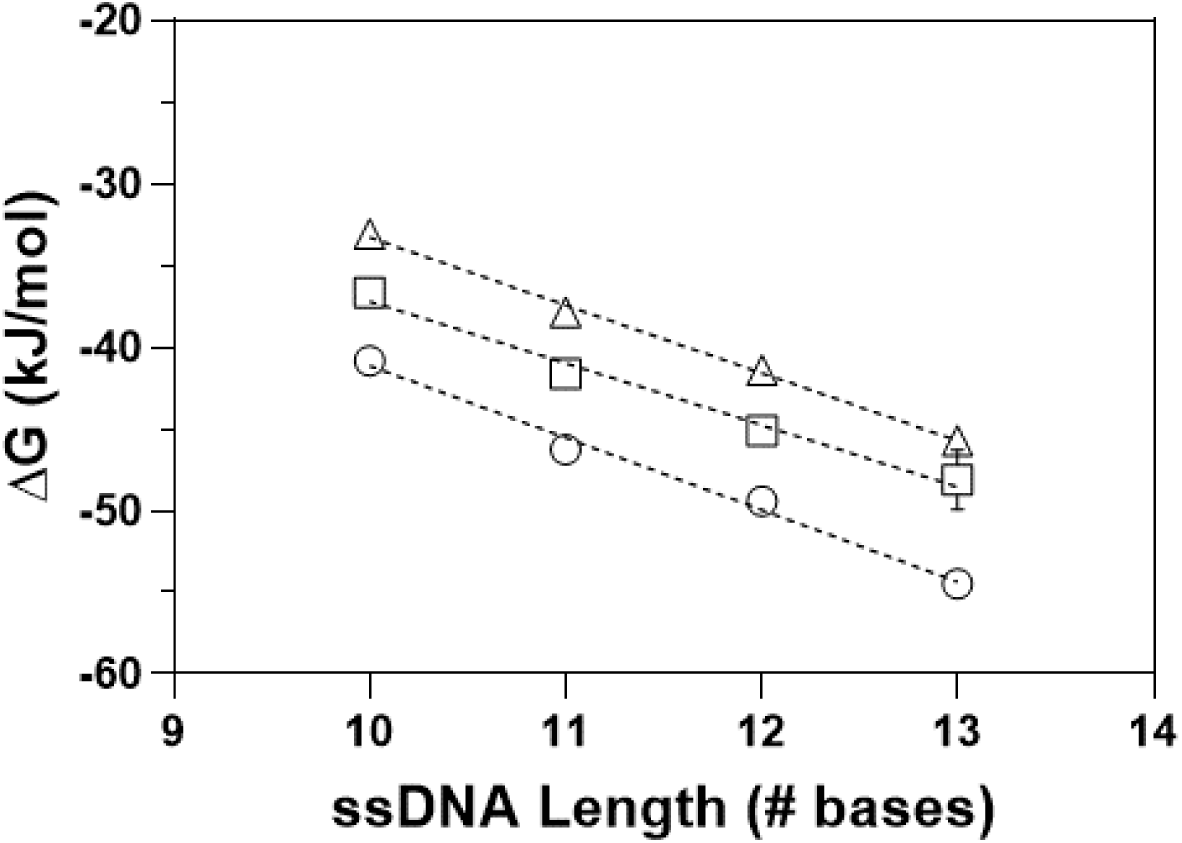
Free energy change for each ASO with respect to ssDNA base length and temperature. Circle: 20 °C, Square: 25 °C, Triangle: 30 °C. Free energy values derived from the kinetic experimental values are presented in Table 3.

### 3. Experimental Thermodynamic Analysis Compared to Nearest Neighbor Predictions

One of the original methods in the past for evaluating the thermodynamic stability of double strand oligonucleotides was to measure the UV absorbance change with increasing temperature to measure a thermal melting curve from which the midpoint of melting (T_m_) could be determined. Breslauer et. al., (1986, 1987) pioneered the use of experimental thermal melting curves to measure the ΔH°, ΔS°, and ΔG°of double strand hybridization for short oligomers (6-16 nt) from which the “nearest-neighbor” algorithm was developed to predict thermodynamic data based on any DNA duplex base pair sequence ^13,19^. Since this time many modified algorithms have been generated using additional duplex thermal melting data and modifications accounting for different solution conditions including different NaCl concentrations^16–18^. The thermal melting technique has been the “gold standard” for measuring thermodynamic parameters for 35 years because it does not require highly technical instrumentation and does not require chemical modifications to the oligo as in fluorescence binding studies. Over the years newer techniques have been developed to measure the hybridization process and estimates of kinetic and thermodynamics of binding. Some of the methods used include: fluorescence-labeled oligonucleotides that measure fluorescence quenching, Förster resonance energy transfer, SPR, isothermal titration calorimetry, and differential scanning calorimetry^29,47–49^. Each of these techniques has their pros and cons for use and the ultimate validation for their application to measure accurate thermodynamic hybridization values is from comparison to the UV-thermal melting method or comparison to the nearest-neighbor predicted thermodynamic values for a specific duplex and buffer condition.

Using the nearest-neighbor model ^14,17,50^ we have calculated the ΔH°, ΔS°, and ΔG° for each AS-10 to AS-13 DNA double strand sequence and compare these predicted values to the kinetic-based experimentally determined values as shown in Table 3. The nearest-neighbor prediction includes NaCl corrections for experiments performed in 50 mM Hepes and 150mM NaCl, pH 7.4. There is excellent comparison between the experimental and predicted values for the ΔG° where for example AS-10 has experimental and predicted values at 20 °C of −40.8 kJ/mol and −41 kJ/mol, respectively and AS-13 has values of −54 kJ/mol and −57 kJ/mol, respectively. A comparison of the experimental and nearest neighbor predicted ΔH° (Figure 5, Table 3) shows that the predicted values are only ~10-20% higher than the experimental values. Further comparisons are useful by comparing the SantaLucia nearest-neighbor model predicted ΔH° values for each of the 10 base pairs to the kinetic experimentally derived ΔH° per nearest neighbor base pair. The SantaLucia model predicts a range of base pair-base pair ΔH° energetics with −31.2 kJ/mol for AA/TT pairs (as the lowest) and −43.46 kJ/mol for CG/GC pairs (as the highest)^17^. The experimentally observed average ΔH° per nearest neighbor base pair for AS-10 (9 pairs) is −29.7 kJ/mol which is unsurprisingly also only different by ~ 10-20% of the nearest-neighbor model. Finally, both experimental and nearest-neighbor methods agree that the driving force for hybridization is a more favorable exothermic enthalpy of binding than entropic.

**Figure 5.**
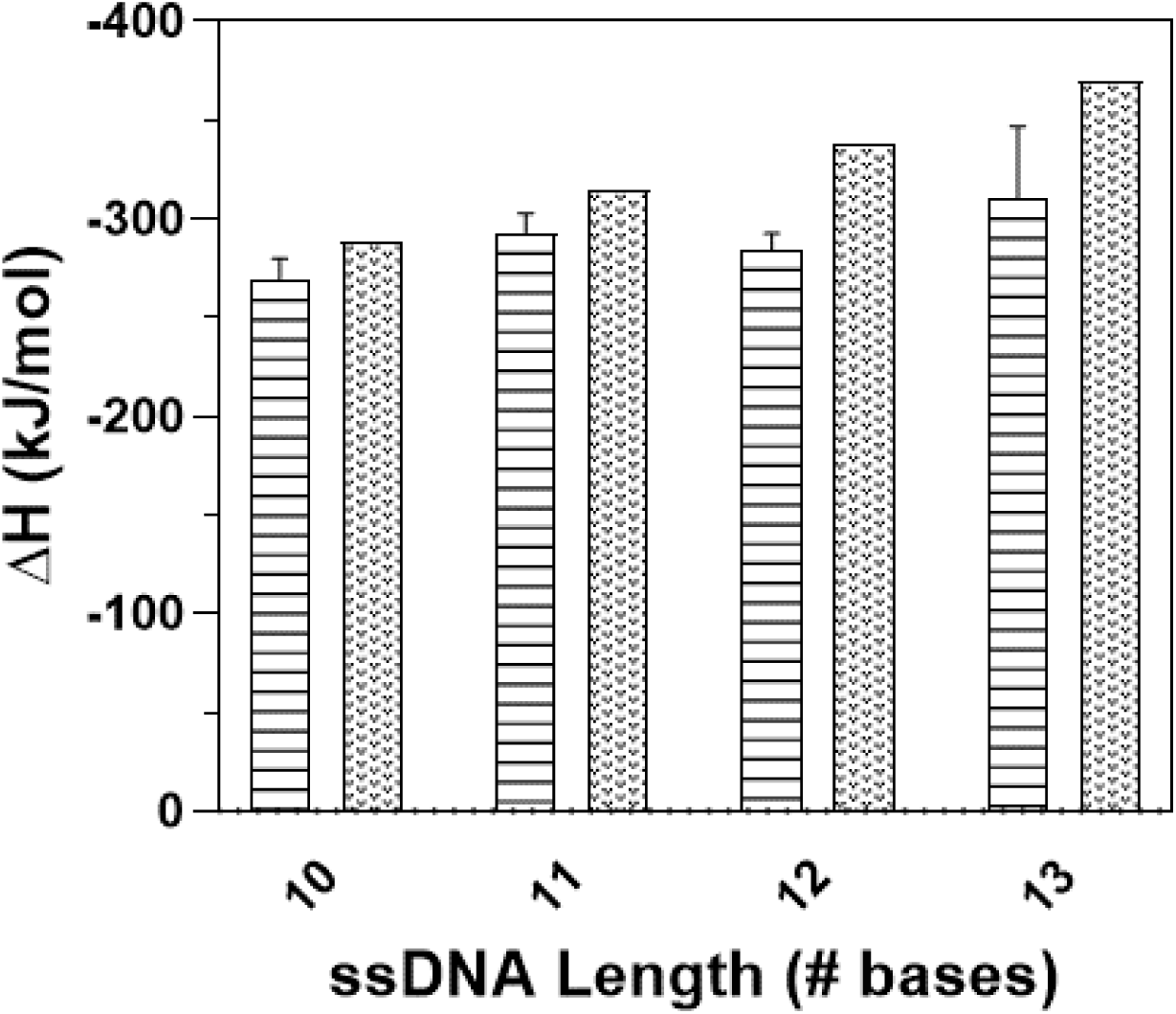
Comparison of the kinetic experimentally derived ΔH° to the Nearest Neighbor model prediction. The AS-10, AS-11, AS-12 and AS-13 data in the horizontal hash bars were derived from Table 3. The Nearest Neighbor predicted ΔH° are the spotted bars.

The thermodynamic information generated for these DNA antisense oligomers binding to the synthetic DNA HIF1α sense Intron-2-Exon-3 “3’splice site” is important for providing guidance on what the affinity might be for longer ASO and affinity at the more physiological temperature 37 °C. This information is also useful as control data for the HIF1α PNA antisense oligomers described by Chung et. al., (2018)^33^ which explored activity of oligomers 10-26 nucleotide long in cell-based assays and *in vivo* models. Since the nearest-neighbor model predicted the ΔG° very well we have used it to provide information on what the affinity (K_d_) would be for longer HIF1α antisense duplexes at 37 °C. The predicted nearest neighbor K_d_ at 37 °C (175 mM Na^+^) for AS-13 is 276 nM. The predicted K_d_ for the corresponding HIF1α 15mer and 17mer antisense strands would be 9.8 nM and 1.1 nM at 37 °C. This calculation makes it clear that the affinity of DNA single strand ASO decreases greatly at physiological temperature (37 °C) and that is one reason why DNA antisense lengths of 17 nucleotides or longer may be important for robust cellular response. The importance of having long oligonucleotides sequences to increase affinity becomes less important when ASO are designed with a modified backbone or nucleotides such as PNA. Modified ASO such as PNAs have higher intrinsic hybridization affinity due to factors such as the absence of the inter-strand electrostatic repulsion from the two negatively charged phosphate backbones. Estimates of the affinity for a HIF1α DNA 15mer and 17mer ASO at 37 °C also enables calculations of the effect on the dissociation rate constant, k_d_, and dissociation half-life (t_1/2_) which are factors determining the time that the ASO is bound to the intracellular target sense strand. As mentioned above the nearest-neighbor predicted K_d_ is 9.8 nM and 1.1 nM (15-mer and 17-mer, 37 °C) and assuming a k_a_ of 4.0 10^5^ M^−1^s^−1^ (based on SPR k_a_ at 30 °C), which is likely to be fairly invariant with different short DNA oligomers, then the dissociation halftimes, t_1/2_, are 2.9 min and 26 min for the ssDNA 15mer and 17mer, respectively^44,51^. These high rates of dissociation may not be desirable for an ASO drug, and this is an example of how DNA ASO kinetic and thermodynamic studies as presented here can provide guidance on optimizing both kinetic and affinity properties in a preclinical setting as well as justify developing PNA-like antisense drugs.

## 4. Conclusions

The technology involved in developing therapeutic antisense drugs is relatively new and requires a wide range of novel synthesis and analytical characterizations to identify potential clinical candidates. The use of SPR to measure kinetic and thermodynamic properties of oligonucleotide hybridization is not new, however as we describe above, the experimental design and analysis of some early studies were not optimal. In this study we demonstrate how to perform a systematic hybridization kinetic and thermodynamic analysis. We used HIF1α ssDNA antisense strands to provide baseline information that would guide our understanding of future PNA-based HIF1α Intron2-Exon3 “3’-splice site” antisense drugs. Examples of some of the fundamental results that can be obtained by SPR kinetic studies include: determining that the affinity (K_d_) for the AS-10 10mer is relatively weak (K_d_, 54 nM) but increases greatly (270-fold) by adding only 3 more HIF1α base pairs resulting in a K_d_ of 0.2 nM for AS-13 at 20 °C. The kinetic analysis for the hybridization makes it clear that the increase in affinity is driven nearly completely by a 150-fold decrease in the dissociation off rate, k_d_, (AS-10 to AS-13) or an ~ 5-fold decrease per each added base pair. Thus, as has been reported before, the design of potent antisense strands is favored by nucleotide length which may also improve selectivity.

Performing temperature dependent SPR kinetic studies enabled the calculation of the enthalpy, entropy and free energy of binding using the van’t Hoff plot which can be used to estimate the energetics of hybridization for longer antisense strands and at physiological temperature. The kinetic studies indicate the ΔH° for the AS-10 hybridization is −267 kJ/mol indicating the base pair hydrogen bond and stacking energetics is ~ −29 kJ/mol per nearest neighbor pair (9 for AS-10) for a nucleotide sequence having 6 A:T and 4 G:C base pairs. The overall ΔG° of stabilization for AS-10 is −33 kJ/mol or ~ −3.7 kJ/mol per nearest neighbor pair at 30 °C. The excellent comparison between the kinetic temperature-dependent thermodynamics (ΔH°, ΔS°, ΔG°) of hybridization and that derived from a nearest-neighbor prediction (Table 3) supports the use of the nearest-neighbor method to predict what the thermodynamic properties would be for HIF1α antisense strands longer than 13 bases at the physiological temperature of 37 °C. The experimentally extrapolated K_d_ for the AS-13 13mer at 37 °C would be very weak at ~ 270 nM. This type of information is useful to identify properties to improve including estimating antisense lengths that would have a more desirable affinity for the development of a drug at 37 °C. Using the nearest-neighbor prediction an HIF1α ASO containing 17 nucleotides spanning the Intron2-Exon3 splice site would have a K_d_ of ~ 1 nM which is on the order of an affinity likely required for an antisense drug. These examples show how a carefully designed SPR temperature-dependent kinetic analysis can provide critical thermodynamic and mechanism of action information for designing antisense drugs.

## Funding

This work was supported by Olipass Corporation by a research grant (ID 2103024) provided to Stevens Institute of Technology.

## Acknowledgements

We would like to thank the Olipass Corporation for their support of this project. We would also like to thank Peter Tolias for his support and helpful discussions.

## Notes

### Competing Interest Statement

The authors have declared no competing interest.

### Summary of Updates

Inadvertent typographical errors were identified in the abstract. These have been corrected to clarify the presentation of the results.

